# MRCA time and epidemic dynamics of the 2019 novel coronavirus

**DOI:** 10.1101/2020.01.25.919688

**Authors:** Chi Zhang, Mei Wang

## Abstract

The 2019 novel coronavirus (2019-nCoV) have emerged from Wuhan, China. Studying the epidemic dynamics is crucial for further surveillance and control of the outbreak. We employed a Bayesian framework to infer the time-calibrated phylogeny and the epidemic dynamics represented by the effective reproductive number (*R*_*e*_) changing over time from 33 genomic sequences available from GISAID. The time of the most recent common ancestor (MRCA) was December 17, 2019 (95% HPD: December 7, 2019 – December 23, 2019). The median estimate of *R*_*e*_ shifted from 1.6 to 1.1 on around January 1, 2020. This study provides an early insight of the 2019-nCoV epidemic. However, due to limited amount of data, one should be cautious when interpreting the results at this stage.

## Introduction

An outbreak of a novel coronavirus (2019-nCoV) was reported in Wuhan, a city in central China (WHO). Coronaviruses cause diseases range from common cold to severe pneumonia. Two fatal coronavirus epidemics over the last two decades were severe acute respiratory syndrome (SARS) in 2003 and Middle East respiratory syndrome (MERS) in 2012 (WHO). Human to human transmission has been confirmed for this new type of coronavirus (Wang et al. 2020) and more than 8,000 cases have been reported as of January 29, 2020.

Studying the virus epidemic dynamics is crucial for further surveillance and control of the outbreak. Phylogeny of the viruses is a proxy of the transmission chain. In this study, we used the birth-death skyline serial (BDSS) model (Stadler et al. 2013) to infer the phylogeny, divergence times and epidemic dynamics of 2019-nCoV. This approach takes the genomic sequences and sampling times of the viruses as input, and co-estimates the phylogeny and key epidemic parameters in a Bayesian framework while accounting for their uncertainties. Particularly, we estimated the shifting time and values of the effective reproductive number (*R*_*e*_) to detect the effect of the intervention.

## Results and Discussion

The sources of the genomic sequences are given in Table 1. The phylogeny in Figure 1 shows the divergence times and relationships of the 33 BetaCoV viruses. Note that this phylogeny is a maximum clade credibility (MCC) tree summarized from the posterior samples, which represents a best estimate of the topology. Due to the similarity of the sequences, the probabilities in most clades are very low (< 0.5) and would form polytomies if summarized as a 50% majority-rule consensus tree (GISAID). The epidemic parameters were estimated while taking the topological uncertainties into account by averaging them in the Bayesian Markov chain Monte Carlo (MCMC) algorithm.

**Table 1.** Data from GISAID.

| Virus name | Accession ID | Collection date |
| --- | --- | --- |
| BetaCoV/Wuhan/IVDC-HB-01/2019 | EPI_ISL_402119 | 2019/12/30 |
| BetaCoV/Wuhan/IVDC-HB-04/2020 | EPI_ISL_402120 | 2020/1/1 |
| BetaCoV/Wuhan/IVDC-HB-05/2019 | EPI_ISL_402121 | 2019/12/30 |
| BetaCoV/Wuhan/IPBCAMS-WH-01/2019 | EPI_ISL_402123 | 2019/12/24 |
| BetaCoV/Wuhan/WIV04/2019 | EPI_ISL_402124 | 2019/12/30 |
| BetaCoV/Wuhan-Hu-1/2019 | EPI_ISL_402125 | 2019/12/26 |
| BetaCoV/Wuhan/WIV02/2019 | EPI_ISL_402127 | 2019/12/30 |
| BetaCoV/Wuhan/WIV05/2019 | EPI_ISL_402128 | 2019/12/30 |
| BetaCoV/Wuhan/WIV06/2019 | EPI_ISL_402129 | 2019/12/30 |
| BetaCoV/Wuhan/WIV07/2019 | EPI_ISL_402130 | 2019/12/30 |
| BetaCoV/Wuhan/HBCDC-HB-01/2019 | EPI_ISL_402132 | 2019/12/30 |
| BetaCoV/Wuhan/IPBCAMS-WH-04/2019 | EPI_ISL_403929 | 2019/12/30 |
| BetaCoV/Wuhan/IPBCAMS-WH-03/2019 | EPI_ISL_403930 | 2019/12/30 |
| BetaCoV/Wuhan/IPBCAMS-WH-02/2019 | EPI_ISL_403931 | 2019/12/30 |
| BetaCoV/Guangdong/20SF012/2020 | EPI_ISL_403932 | 2020/1/14 |
| BetaCoV/Guangdong/20SF013/2020 | EPI_ISL_403933 | 2020/1/15 |
| BetaCoV/Guangdong/20SF014/2020 | EPI_ISL_403934 | 2020/1/15 |
| BetaCoV/Guangdong/20SF025/2020 | EPI_ISL_403935 | 2020/1/15 |
| BetaCoV/Guangdong/20SF028/2020 | EPI_ISL_403936 | 2020/1/17 |
| BetaCoV/Guangdong/20SF040/2020 | EPI_ISL_403937 | 2020/1/18 |
| BetaCoV/Nonthaburi/61/2020 | EPI_ISL_403962 | 2020/1/8 |
| BetaCoV/Nonthaburi/74/2020 | EPI_ISL_403963 | 2020/1/13 |
| BetaCoV/Zhejiang/WZ-01/2020 | EPI_ISL_404227 | 2020/1/16 |
| BetaCoV/Zhejiang/WZ-02/2020 | EPI_ISL_404228 | 2020/1/17 |
| BetaCoV/USA/IL1/2020 | EPI_ISL_404253 | 2020/1/21 |
| BetaCoV/USA/WA1/2020 | EPI_ISL_404895 | 2020/1/19 |
| BetaCoV/Taiwan/2/2020 | EPI_ISL_406031 | 2020/1/23 |
| BetaCoV/USA/CA1/2020 | EPI_ISL_406034 | 2020/1/23 |
| BetaCoV/USA/CA2/2020 | EPI_ISL_406036 | 2020/1/22 |
| BetaCoV/USA/AZ1/2020 | EPI_ISL_406223 | 2020/1/22 |
| BetaCov/Wuhan/WH01/2019 | EPI_ISL_406798 | 2019/12/26 |
| BetaCov/Wuhan/WH03/2020 | EPI_ISL_406800 | 2020/1/1 |
| BetaCov/Wuhan/WH04/2020 | EPI_ISL_406801 | 2020/1/5 |

**Figure 1.**
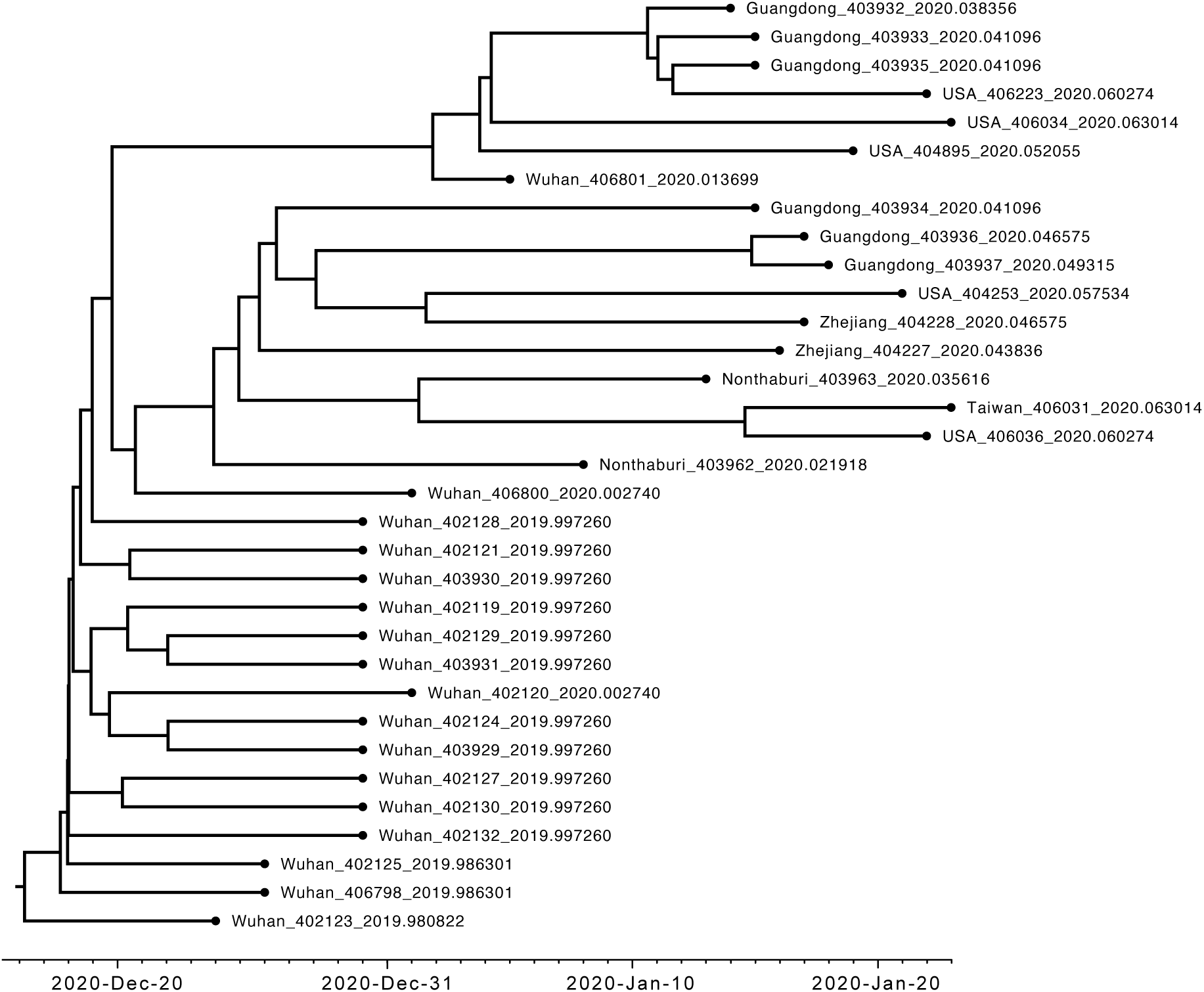
Maximum clade credibility (MCC) tree summarized from the MCMC sample. The common ancestor heights were used to annotate the clade ages.

The time of the most recent common ancestor (MRCA) is estimated to be December 17, 2019 (95% HPD: December 7, 2019 – December 23, 2019) (Table 2). This is in agreement with the symptom onset reported by WHO and several preliminary studies (http://virological.org). The origin time estimated is just a couple of days older than the MRCA time. It appears too young and likely due to unsampled cases not included in our analysis (du Plessis and Pybus 2020).

**Table 2.** Posterior estimates of key model parameters.

|  | median and 95% HPD interval |
| --- | --- |
| $t_0$ | 0.1089 (0.0871, 0.1505) |
| $t_{\text{mrca}}$ | 0.1005 (0.0852, 0.1284) |
| $R_{e1}$ | 1.57 (0.78, 3.64) |
| $R_{e2}$ | 1.13 (0.67, 1.81) |
| $t_{\text{shift}}$ | 2020.0020 (2019.9830, 2020.0311) |
| $\delta$ | 84.91 (18.37, 187.64) |
| $p$ | 0.19 (0.019, 0.43) |
| $r$ | 0.0016 (0.00076, 0.0027) |
| $\kappa$ | 5.05 (2.05, 9.34) |
| $\alpha$ | 0.67 (0.0011, 3.03) |

Note: time unit is years.

We investigate the epidemic dynamics of 2019-nCoV by estimating *R*_*e*_ before and after a shifting time. *R*_*e*_ > 1.0 means that the number of cases are increasing and the epidemic is growing, whereas *R*_*e*_ < 1.0 means that the epidemic is declining and will die out. Interestingly, the median estimate of *R*_*e*_ shifted from 1.6 to 1.1 on around January 1, 2020 (Table 2). In general, this is in agreement with some other studies reporting *R*_*e*_ ranging from 1.4 to 5.5 (Read et al. 2020; Zhao et al. 2020; Riou and Althaus 2020) and the intervention happened around January 1 (Li et al. 2020).

Keep in mind that we used only 33 samples in our analysis, which is less than 1% of the reported number of infected patients, thus one needs to be cautious when interpreting the results. With more viruses sequenced, we would expect more reliable estimates which would provide better insights into the epidemic of 2019-nCoV.

Overall, this study provides an early insight of the 2019-nCoV epidemic by inferring key epidemiological parameters from the virus sequences. Such estimates would help public health officials to coordinate effectively to control the outbreak.

## Material and Methods

We collected 33 genomic sequences available from GISAID (Table 1). Sequences were aligned using MUSCLE (Edgar 2004). The first and last 150bp were removed, resulting in a total length of 29604bp for the alignment. The collection dates of the viruses ranged from December 24, 2019 to January 23, 2020 and they were used as fixed ages (in unit of years) in subsequent analysis.

We used the BDSS model (Stadler et al. 2013) implemented in the BDSKY package for BEAST 2 (Bouckaert et al. 2019) to infer the phylogeny, divergence times and epidemic dynamics of 2019-nCoV. The model has an important epidemiological parameter, the effective reproductive number *R*_*e*_, defined as the number of expected secondary infections caused by an infected individual during the epidemic. The model allows *R*_*e*_ to change over time, making it feasible to estimate its dynamics (Stadler et al. 2013). In our case, we just allowed one shift of *R*_*e*_ at time *t*_shift_ and co-estimated them. The prior for *R*_*e*_ was a lognormal(0, 1.25) distribution with median 1.0 and 95% quantiles between 0.13 and 7.82, and that for *t*_shift_ was normal(2020.010959, 0.010959) with mean on January 4 and standard deviation of 4 days. The BDSS process starts from the origin time *t*_0_, which was assigned a lognormal(–1, 1.5) prior with median 0.368 (years before the latest sampling time). The other two parameters are the becoming noninfectious rate *δ* and sampling proportion *p*, which were assumed constant over time. *δ* was given a lognormal (2, 1.25) prior with median 7.39 and mean 16.1, expecting the infectious period of an individual (1/*δ*) to be about a month. The sampling proportion of infected individuals *p* was a beta(1, 9) distribution with mean 0.1.

We assumed a strict clock and the clock rate *r* was assigned a gamma(2, 0.0005) prior with mean of 0.001 substitutions per site per year. The substitution model used was HKY+Γ_4_ (Hasegawa et al. 1985; Yang 1994) in which the transition-transversion rate ratio *κ* was set a lognormal(1, 1.25) prior and the gamma shape parameter *α* was an exponential(1) prior.

The analysis was performed in the BEAST 2 platform (Bouckaert et al. 2019). We ran 100 million MCMC iterations and sampled every 5000 iterations. The first 20% samples were discarded as burn-in. Convergence was diagnosed in Tracer (Rambaut et al. 2018) to confirm that independent runs gave consensus results and all parameters had effective sample size (ESS) larger than 100. The remaining 80% samples were used to build the maximum clade credibility (MCC) tree and to summarize the parameter estimates.

## Acknowledgments

We thank Louis du Plessis for his valuable help and suggestions on the analysis.

